# Whole blood metabolomics of dementia patients reveal classes of disease-linked metabolites

**DOI:** 10.1101/2021.06.22.449525

**Authors:** Takayuki Teruya, Yung-Ju Chen, Hiroshi Kondoh, Yasuhide Fukuji, Mitsuhiro Yanagida

**Affiliations:** G0 Cell Unit, Okinawa Institute of Science and Technology Graduate University (OIST), 1919-1 Tancha, Onna-son, Okinawa, 904-0495 Japan; Geriatric unit, Graduate School of Medicine, Kyoto University, Sakyo-ku, Kyoto 606-8507, Japan; National Hospital Organization Ryukyu Hospital, 7958-1 Kin, Kin-cho, Okinawa 904-1201, Japan

**Keywords:** dementia, Alzheimer’s disease, antioxidants, ergothioneine, trimethyl-ammonium compounds

## Abstract

Dementia is caused by factors that damage neurons. We quantified small molecular markers in whole blood of dementia patients, using non-targeted liquid chromatography-mass spectroscopy (LC-MS). Thirty-three metabolites, classified into 5 groups (A-E), differed significantly in dementia patients, compared with healthy elderly subjects. Seven Group A metabolites present in plasma, including quinolinic acid, kynurenine, and indoxyl-sulfate, increased. Possibly they act as neurotoxins in the central nervous system (CNS). The remaining 26 compounds (Groups B-E) decreased, possibly causing a loss of support or protection of the brain in dementia. Six Group B metabolites, normally enriched in red blood cells (RBCs) of healthy subjects, all contain trimethylated ammonium moieties. These metabolites include ergothioneine and structurally related compounds have scarcely been investigated as dementia markers, validating the examination of RBC metabolites. Ergothioneine, a potent anti-oxidant, is significantly decreased in various cognition-related disorders, such as mild cognitive impairment and frailty. Group C compounds, also include some oxidoreductants and are normally abundant in RBCs (NADP^+^, glutathione, ATP, pantothenate, S-adenosyl-methionine, and gluconate). Their decreased levels in dementia patients may also contribute to depressed brain function. Groups D (12) contains plasma compounds, such as amino acids, glycerophosphocholine, dodecanoyl-carnitine, 2-hydroxybutyrate, which normally protect the brain, but their diminution in dementia may reduce that protection. Seven Group D compounds have been identified previously as dementia markers. Group B-E compounds may be critical to maintain the CNS by acting directly or indirectly. How RBC metabolites act in the CNS and why they diminish so significantly in dementia remain to be determined.

**Significance Statement:** Dementia is a slowly progressing, chronic, and usually irreversible decline in cognitive function. Mechanistic causes and definitive treatments remain elusive. Using comprehensive metabolomics, we identified 5 groups of metabolites (A-E), 21 of which are novel, possibly useful for diagnosis and therapy of forms of dementia, such as Alzheimer’s disease. Seven Group A compounds may act as neurotoxins, whereas Group B-E compounds may protect the CNS against oxidative stress, maintain energy reserves, supply nutrients and neuroprotective factors. Five metabolites, ergothioneine, *S*-methyl-ergothioneine, trimethyl-histidine, methionine, and tryptophan identified in this study overlap with those reported for frailty. Interventions for cognitive diseases involving these dementia metabolomic markers may be accomplished either by inhibiting Group A compounds or by supplementing Group B-E compounds in patients.

## Introduction

Dementia is a collective term to describe various symptoms of cognitive impairment in a condition in which intelligence is irreversibly diminished due to acquired organic disorders of the brain, characterized by deterioration of memory, thinking, behavior, and the ability to perform daily activities (1, 2). Though a common cause is Alzheimer’s disease (AD), a neurodegenerative disease in which memory is rapidly impaired due to hippocampal atrophy, multiple types of dementia, known as mixed dementia, can coexist (3, 4). Mental and physical exercise and avoidance of obesity may reduce the risk of dementia (5-7). No medications or supplements have been definitively shown to decrease risk (8, 9). Dementia most often begins in people over 65 years of age, and about 6% of seniors are afflicted with it. It is one of the most costly diseases in developed countries (10).

In this study, we conducted non-targeted, comprehensive analysis of blood metabolites in dementia patients. Thorough metabolomic evaluation can supply complete information about metabolite abundance in each subject. While non-targeted analysis is far more laborious than targeted analysis, the effort expended in this “no assumptions” approach is often recompensed by identification of diagnostic compounds overlooked by targeted analysis. A wealth of metabolite information may provide clues to understanding the profound metabolic changes occurring in dementia. Liquid chromatography-mass spectroscopy (LC-MS) was employed for whole blood metabolite profiling of dementia patients, and we found metabolic compounds not previously known to be related to dementia. Metabolomics of blood cells have scarcely been investigated, particularly in relation to diseases, despite the fact that red blood cells (RBCs) account for about 40% of all blood metabolites (11-13). Thus, metabolomic information from RBCs also provides crucial information on health and disease (14-16). Here we identified 33 dementia-linked markers (12 of which are RBC-enriched) and validated them by principal component analysis (PCA), correlation, and heatmap analyses, confirming that these markers actually are involved in development of dementia. Our results suggest that detailed molecular diagnosis of dementia is now possible. Somewhat unexpectedly, markers deduced from dementia only partially overlap with amino acid markers obtained from frailty patients with cognitive defects (16), so that frailty and dementia partly share the diminished cognitive markers. We also show that an antioxidant ergothioneine, an RBC component involved in human cognitive ability (16, 17) and two related compounds are reduced in dementia.

## Results

### Collection of blood samples from dementia subjects

To identify dementia-related blood metabolites, quantitative comparisons were conducted of blood samples of dementia patients, healthy elderly (HE), and healthy young (HY) subjects. Blood samples of the dementia patients (age, 75∼88 y) diagnosed and hospitalized at the National Hospital Organization Ryukyu Hospital, Kin-town, Okinawa were obtained from each patient after informed consent (see Materials and Methods). The same number of HE (67∼80 y) and HY (28∼34 y) volunteers from Onna Clinic, Onna-village, Okinawa were also recruited (*SI Appendix*, Table S1 and Fig. S1). Twenty-four subjects comprising 8 dementia patients, 8 healthy elderly (HE), and 8 healthy young (HY) subjects participated in this study. All blood samples were drawn at each hospital, as described (14). Venous blood samples were taken into tubes with heparin as an anti-coagulant.

### Thirty-three dementia-linked blood metabolites comprise 5 subgroups A-E

In all whole blood samples collected, 124 metabolites were identified and quantified by non-targeted LC-MS (*SI Appendix*, Table S2). They consisted of 14 subgroups. Fifty-one compounds, comprising 5 subgroups (nucleotides, vitamins and coenzymes, nucleotides-sugar derivatives, sugar phosphates, and anti-oxidants) are RBC-enriched (14). Of these 124 compounds, 33 metabolites differed significantly between dementia patients and HE (the range of p-value, 0.00016 < *P* < 0.05) (Table 1 and Fig. 1). Five compounds, ATP, glutathione disulfide, glutamine, phenylalanine, and betaine are highly abundant (ranked H). Five other compounds, glycerophosphocholine, ergothioneine, methionine, tryptophan, and tyrosine are of high to medium (H-M) abundance. Three additional compounds vary widely in abundance among healthy subjects (H-L): caffeine, dimethyl-xanthine, and trimethyl-tryptophan. The remaining 20 compounds are of medium to low abundance (M-L, M, L) (Table 1). Twelve of 33 compounds are RBC-enriched, which has been scarcely reported. Characteristically, 9 dementia-related compounds contain trimethyl ammonium moieties (Table 1).

**Table 1.**
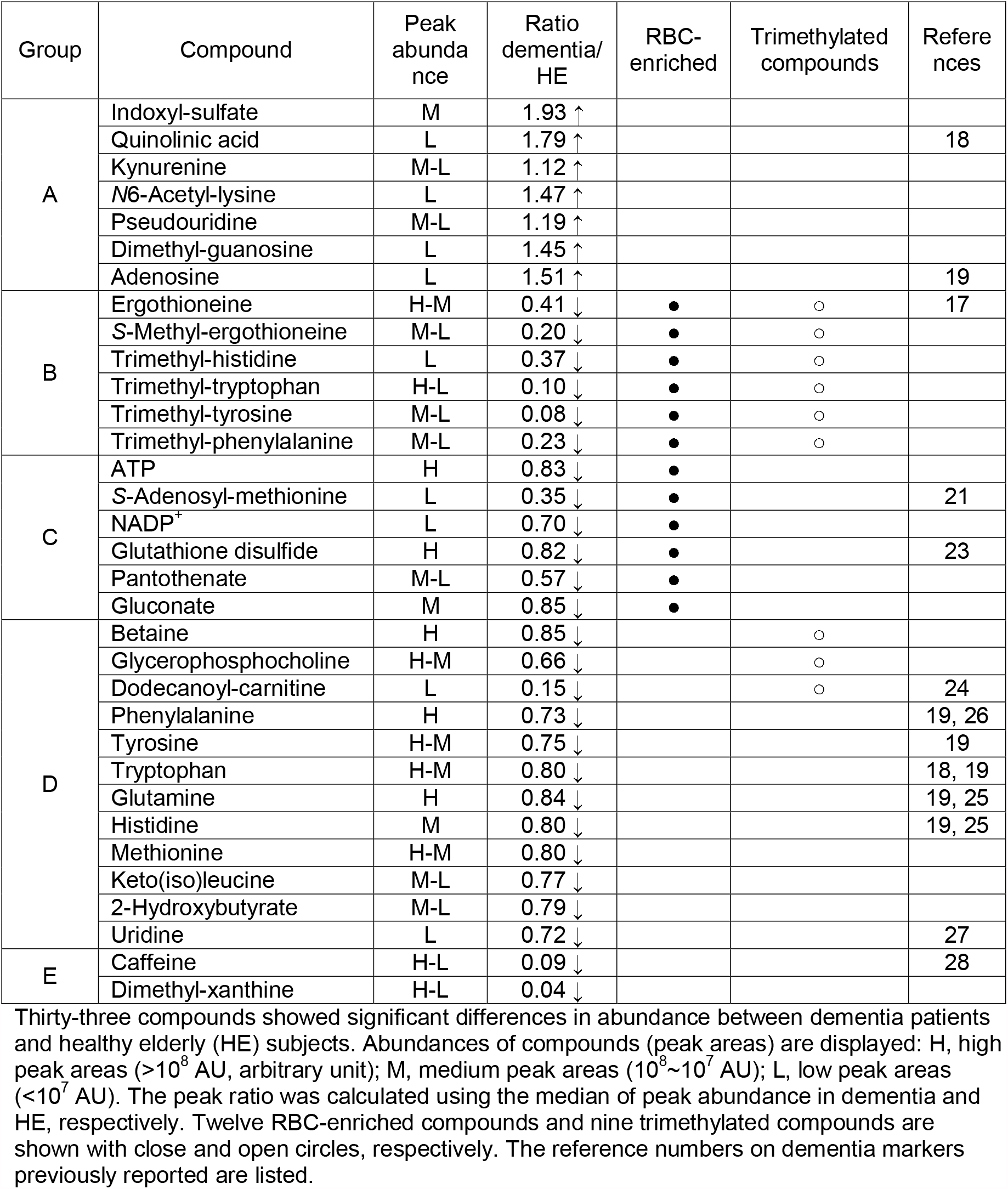
List of 33 dementia markers

**Fig. 1.**
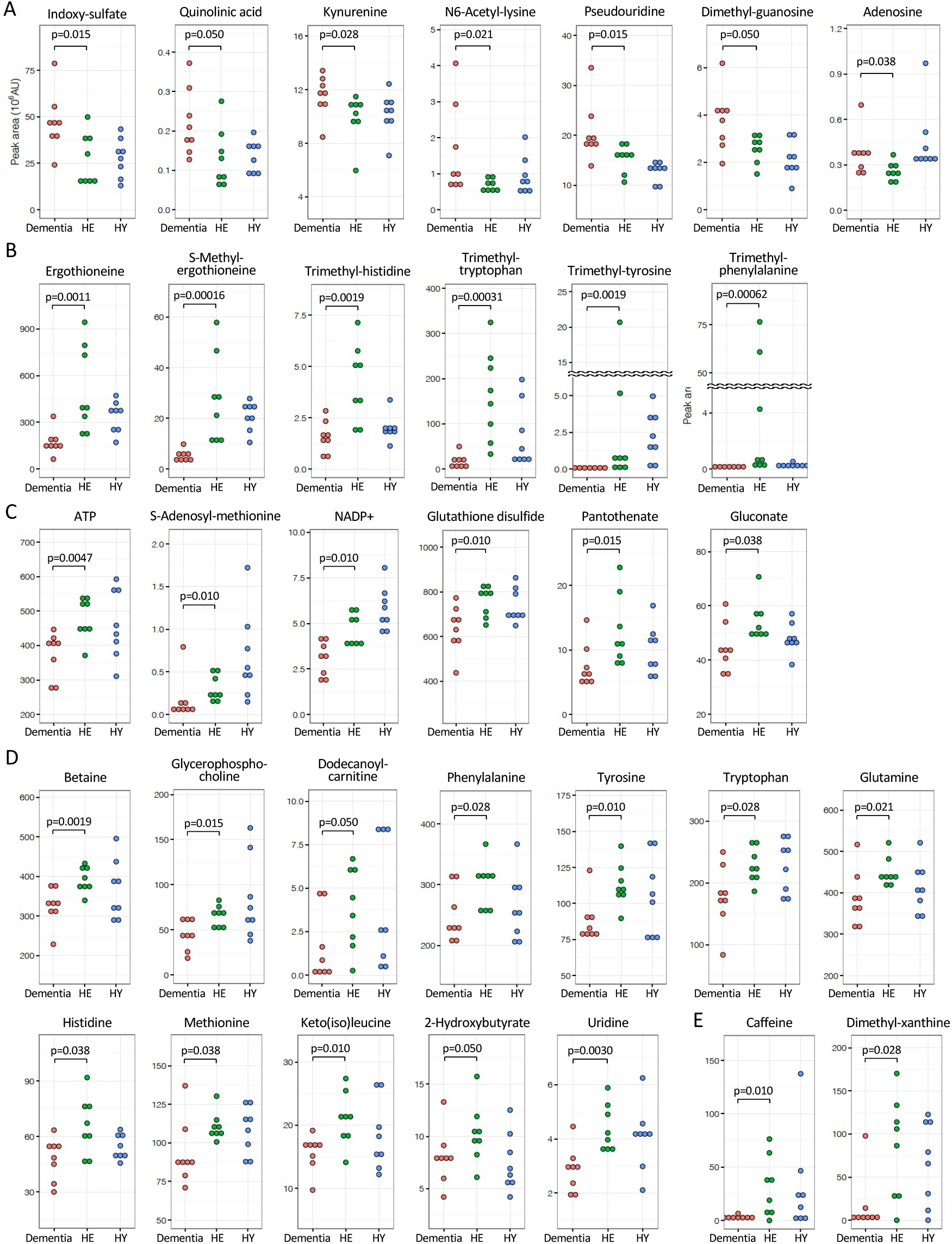
Dot plot profiles of 33 dementia-related metabolites. Dementia metabolites were selected using *P*-values (< 0.05) obtained by comparison of peak abundance between dementia and HE subjects. Compounds were grouped into A-E (see text).

Peak ratios (dementia/HE in Table 1) were calculated using the median of peak abundance in each group. Seven compounds (Group A) that increased in dementia showed peak ratios >1.0. Twenty-six others had ratios <1.0 that declined in dementia. Four compounds exhibiting the greatest decreases are caffeine-related; dimethyl-xanthine (E, 0.04), trimethyl-tyrosine (B, 0.08), caffeine (E, 0.09), and trimethyl-tryptophan (B, 0.10). Curiously, all are aromatics. Their variances are also broad in healthy subjects.

To quantify individual variability of the 124 metabolites, coefficients of variation (CVs) for all experimental populations of 24 subjects were calculated (*SI Appendix*, Table S2). In the 33 dementia-linked compounds, CVs of ATP (0.20), glutathione disulfide (0.14), and NADP^+^ (0.33), which have vital functions, were relatively low, while trimethyl-tryptophan (1.09), trimethyl-tyrosine (2.21), trimethyl-phenylalanine (3.17), and caffeine (1.49) were high. These values were substantially in agreement with those in our previous study, an independent dataset obtained from 30 healthy elderly and young people (14). Thus, the great variability of data in Fig. 1 reflects genuine individual variation in metabolites, which were accurately detected by our metabolomic analysis. These data demonstrate that compounds having small to large individual variability are implicated in dementia.

### Increased Group A compounds may be toxic

Seven Group A compounds were identified by their increases in the dementia patients (Fig. 1*A*). They consisted of indoxyl-sulfate (peak ratio dementia/HE, 1.93), quinolinic acid (1.79), adenosine (1.51), dimethyl-guanosine (1.45), *N*6-acetyl-lysine (1.47), pseudouridine (1.19), and kynurenine (1.12). Two of the 7 metabolites were previously reported as AD-related markers (18, 19). Some of them are reportedly toxic (20), suggesting that they may be inhibitory in brain (see below).

### Twenty-six blood dementia compounds consist of 4 Groups B-E

Twenty-six remaining compounds decreased in dementia patients (*P* < 0.05) (Fig. 1*B–E*). They consisted of 4 subgroups (B-E), having distinct characteristics. Group B compounds include ergothioneine (ET) and 5 other trimethyl-ammonium compounds. To our knowledge, except for ET (17), these are all novel dementia markers, probably because they are enriched in RBCs and scarcely studied in connection with dementia. ET is an anti-oxidant, a thiourea derivative of trimethyl-histidine. Two other ET-related, but less abundant compounds, *S*-methyl-ET and trimethyl-histidine (hercynine), also declined strikingly in blood of dementia patients. A question was addressed if a couple of outliers were eliminated, statistically significant differences would disappear. We therefore attempted to eliminate three of eight subjects with the highest abundances in HE, and found that significant differences (*P* < 0.05) still remained. Thus, our conclusions seem to be valid.

Group C compounds also decreased in dementia patients. These included ATP, NADP^+^ (oxidoreductive coenzyme), glutathione disulfide (GSSG, redox compound), pantothenate (vitamin B5), S-adenosyl-methionine (SAM, methyl donor) (21), and gluconate (zinc carrier) (22). They are related to energy, redox reactions, methylation, and metal ions. Group C compounds were all enriched in RBCs, and four of six are novel dementia markers. Two of them (SAM, GSSG) were previously shown to be AD-related (21, 23).

Trimethyl-tryptophan (hypaphorine), trimethyl-phenylalanine, glycerophosphocholine, dodecanoyl-carnitine (24), and trimethyl-tyrosine, all of which contain trimethyl ammonium ions, also declined. The extent of reduction for trimethyl-tryptophan (0.10) and trimethyl-tyrosine (0.08) was striking. These reductions may be due to instability or reduced synthesis, or to reduced import in dementia patients. Of the nine compounds that contain a trimethyl ammonium moiety, six of them that contain ET are enriched in RBCs and classified as Group B compounds (Table 1).

Twelve Group D metabolites (Table 1) are enriched in blood plasma and seven of them were previously reported to be dementia or AD-markers. They include standard amino acids, glutamine (19, 25), phenylalanine (19, 26), tyrosine (19), histidine (19, 25), methionine, tryptophan (regular amino acids) (18, 19), a pyrimidine nucleoside, uridine (27), and organic acids, 2-hydroxybutyrate (lipid-degradation product) and keto(iso)leucine (keto acid). Caffeine is a known dementia marker (28). Dimethyl-xanthine is a metabolite of caffeine. These greatly declined in dementia and are highly correlated and isolated from other metabolites (see below) so that they are designated Group E. Consistency of Group D plasma metabolites as dementia markers, but not Group B and C RBC metabolites validated the method of searching dementia markers that we employed in the present study. The great majority of metabolites enriched in RBCs were not identified in the previous studies.

### Nine trimethylated ammonium compounds were diminished in dementia patients

Of nine trimethylated compounds that decreased in dementia, six are enriched in RBCs (*SI Appendix*, Fig. S2). Three of them (betaine, glycerophosphocholine, and dodecanoyl-carnitine) are present in plasma and are synthesized in the human body, whereas the other six, containing an aromatic moiety, are derived from food (29, 30). Most strikingly, six of these nine compounds are highly abundant (H, H-M, or H-L) in plasma and RBCs in healthy subjects, and are highly correlated so that their behavior may be highly coordinated. Hence, the sharp declines of these amphipathic compounds (possessing both hydrophilic and lipophilic properties and forming the basis of lipid polymorphism) in blood of dementia patients may strongly affect the physicochemical properties of neuronal systems.

### Seven metabolites increased in dementia

Interestingly, 7 metabolites of Group A comprising 3 nucleosides and 4 amino acid derivatives, increased in dementia (Fig. 1*A*). None was highly abundant and none was enriched in RBCs, and their increase in dementia occurred in plasma. Indoxyl-sulfate, kynurenine, and quinolinic acid (18) are involved in tryptophan metabolism and possibly act as excitatory toxins in brain (31, 32), while *N*6-acetyl-lysine is implicated in histone and non-histone protein modification (33). Pseudouridine, adenosine (19), and dimethyl-guanosine are degradation products of RNAs present in urine and are thought to be oxidized (34, 35). Increases of these metabolites in dementia are of great interest, as some are reportedly toxic in the central nervous system (CNS) and may lead to impairment of the brain (36-38).

### PCA separates dementia patients from non-dementia subjects (HE)

To distinguish between dementia and HE subjects, we then applied principal component analysis (PCA). We calculated principal component (PC1, PC2) values using abundance data of the 33 dementia-related metabolites in 16 subjects. Dementia and HE subjects were clearly separated (Fig. 2*A*). We then attempted to achieve the same degree of resolution using fewer metabolites and found that six metabolites (dimethyl-guanosine, pseudouridine, NADP^+^, trimethyl-histidine, ET, *S*-methyl-ET) were still able to separate dementia from non-dementia subjects almost perfectly (Fig. 2*B*).

**Fig. 2.**
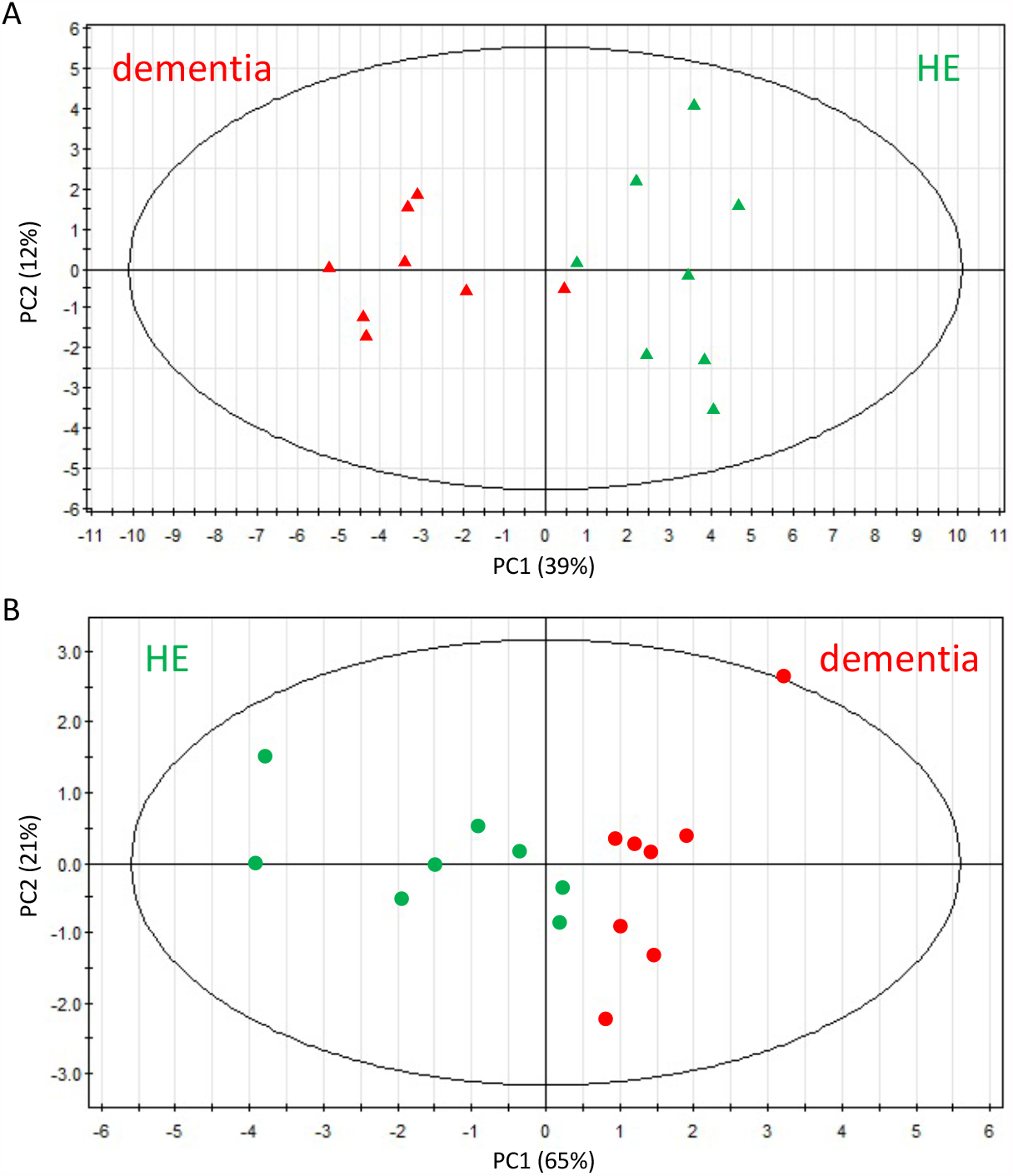
PCA of 33 AD or 6 selected dementia-related compounds showed significant differences between patients with dementia and HE. Blood data of dementia and HE subject were subjected to PCA (principal component analysis). (*A*) A PCA plot using abundances of 33 (7 increased and 26 decreased) dementia-related compounds. Dementia and HE subjects were separated into two domains (see text). Red, dementia; green, HE. (*B*) PCA was also performed using only 6 selected dementia markers (dimethyl-guanosine, pseudouridine, *S*-methyl-ET, ET, trimethyl-histidine, and NADP^+^).

### Correlation analysis corroborates five metabolite groups

Levels of some blood metabolites linked in a biochemical pathway and/or function show correlations (14). We first confirmed correlations within ergothioneine derivatives (ET, S-methyl-ET, trimethyl-histidine) and between caffeine and dimethyl-xanthine using Pearson’s correlation coefficient r. Abundance of these compounds indicated high correlations (0.92 > r > 0.76, p < 0.0001) (*SI Appendix*, Fig. S3) as previously reported (14). Thus, the dataset of the present study is reproducible and statistically valid.

Second, to gain insight into how these 33 dementia metabolites are related, we searched for relationships among them. Significant positive and negative correlations (0.89 > r > 0.50 or - 0.68 < r < -0.50) in Groups A-E support the classification of subgroups shown in Fig. 3.

**Fig. 3.**
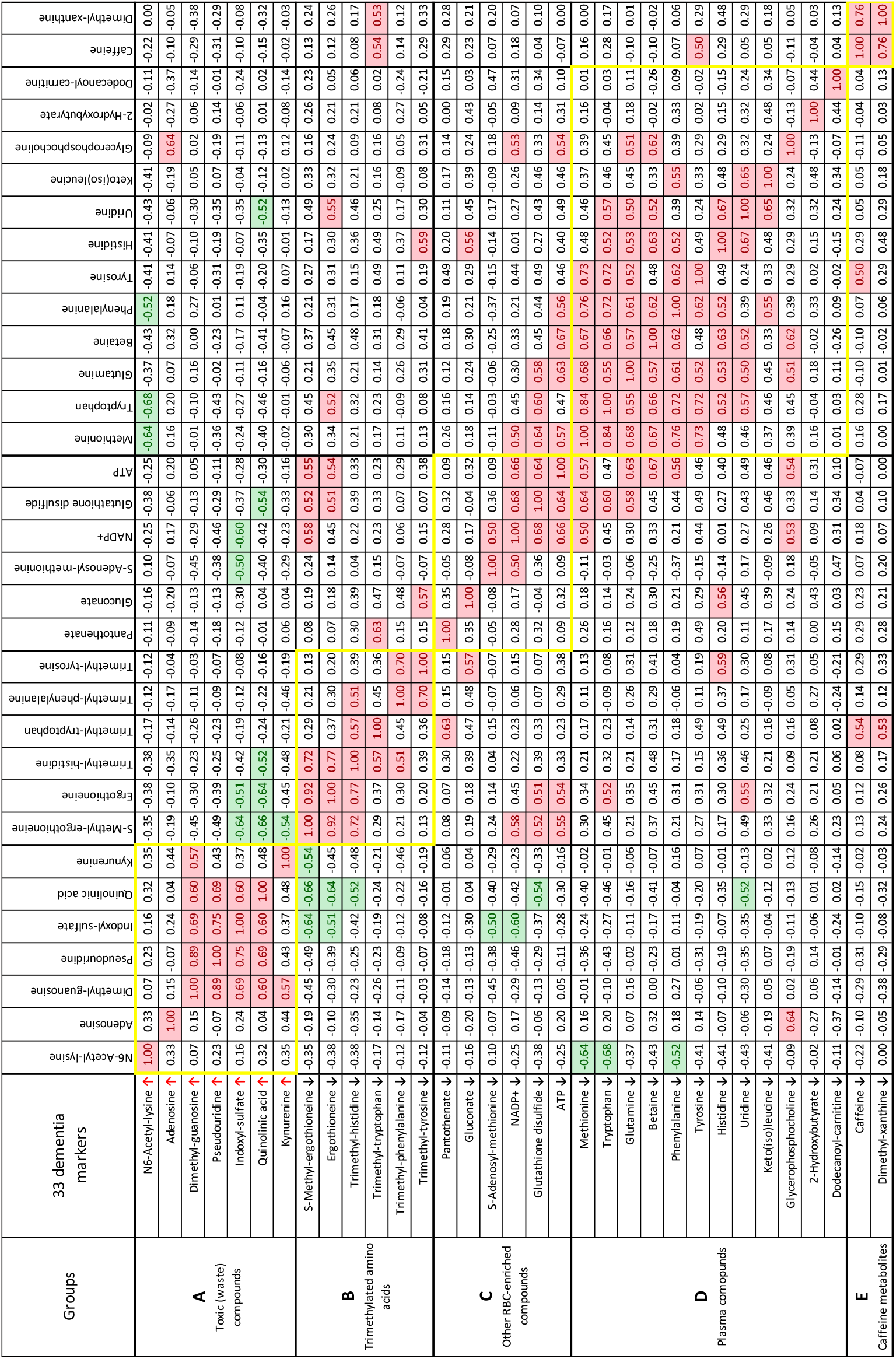
Correlation analysis of 33 dementia markers. Correlation values (*r*) larger than +0.5 are marked with red, whereas values below -0.5 are in green. Compounds that increased or decreased in dementia patients are indicated by arrows. Five subgroups of 33 compounds are shown with frames in yellow.

Five of seven Group A markers were correlated (0.89∼0.57) within the subgroup (quinolinic acid, dimethyl-guanosine, pseudouridine, indoxyl-sulfate, and kynurenine). This striking correlation strongly supported their classification within Group A and oppose to Group B-E. How Group A compounds molecularly oppose to B, C, and D (green color) remains to be determined. The mode of interaction is unknown whether the mode is indirect or direct remains to be studied.

Negative correlations exist not only between Group A and B compounds but also between Group A and C compounds. Group B and C metabolites are correlated within the groups, again validating their subgroup designations. Group B metabolites are structurally related, as they commonly contain trimethylated-ammonium group indicating that they are anti-oxidative. Group C metabolites are not structurally related, but three of them (NADP^+^, glutathione disulfide, and ATP), implicated in redox and energy metabolism, are highly correlated and enriched in RBCs. These distinct metabolites may coordinate to support directly or indirectly their metabolites pathways. Finally, two metabolites, caffeine and dimethyl-xanthine in Group E were highly correlated; the latter is the precursor of caffeine. There are reports on caffeine showing that these purines are beneficial to relieve dementia (see Discussion).

### Heatmap comparison of dementia and HE subjects

Using abundance data of 33 dementia-compounds in 8 dementia patients and 8 HE subjects, a heatmap was constructed (Fig. 4 and *SI Appendix*, Fig. S4). In dementia patients, 7 Group A compounds mostly displayed red color cells in dementia patients due to the increased abundances, whereas HE (healthy elderly) subjects mostly showed the blue color cells due to the decreased abundances. In the remaining 26 compounds, dementia subjects displayed the blue cells, due to their decrease whereas HE subjects showed the red cells due to their increase more than the average. Strikingly, the seven Group A compounds increased nearly uniformly in dementia patients, whereas in HE and also HY (*SI Appendix*, Fig. S4) subjects, the levels were mostly below 50, indicating that these may be appropriate as diagnostic markers for dementia. Thus the heatmap profiles showed individual variations of dementia-linked metabolites, and further study will provide more evidence for the use as a diagnostic tool.

**Fig. 4.**
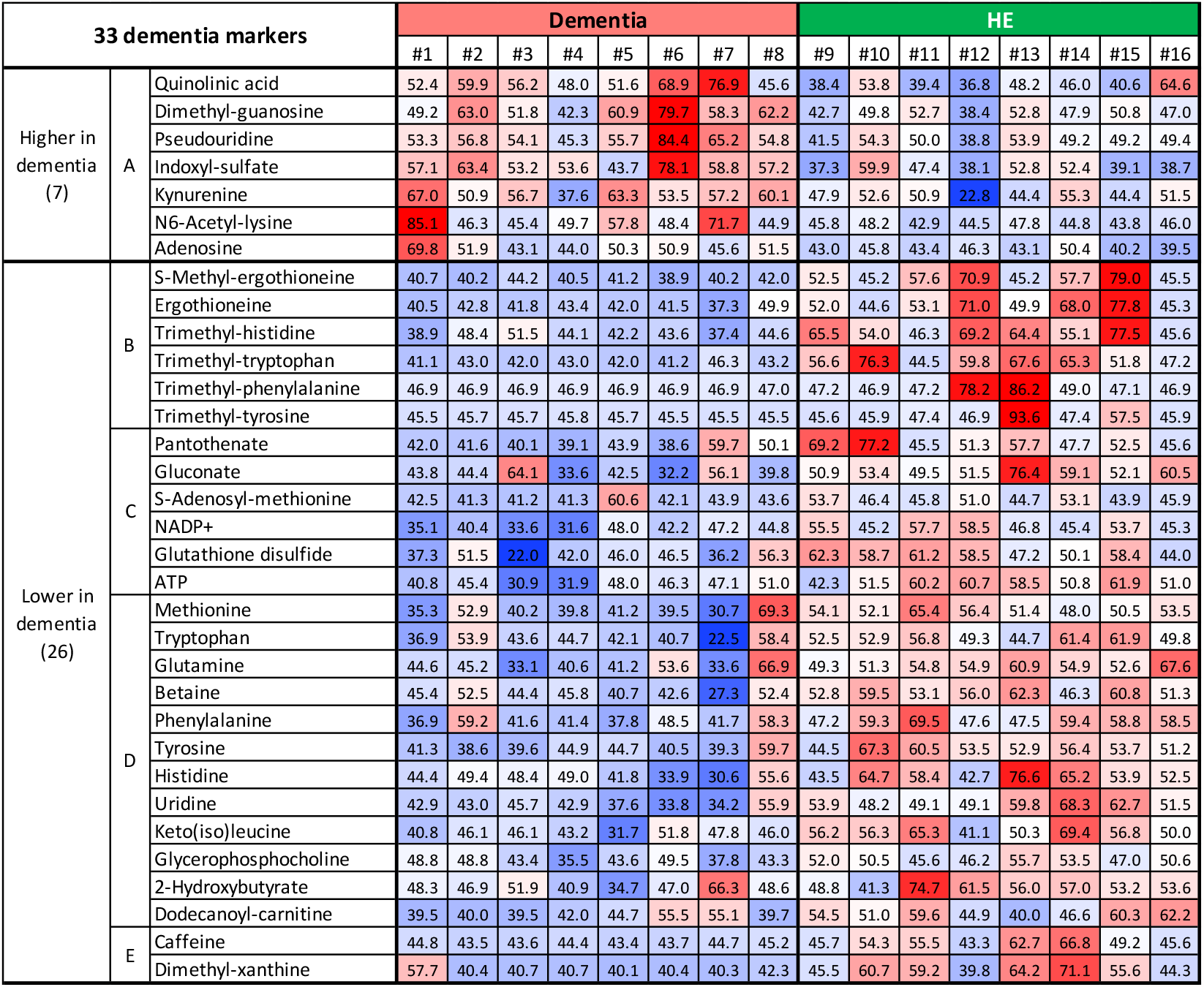
Construction of a heatmap using abundance data of 33 compounds for each subject. Standardized scores (T-scores) are represented by colors. The average value (50), white; values above average, pale red to deep red; values below average, pale blue to deep blue. Seven compounds that increased in dementia tended to be red in dementia whereas they tended to be white to blue in HE. For the 26 remaining compounds, colors are reversed in dementia and HE.

## Discussion

A principal conclusion of this study is that plasma-enriched dementia factors Group A compounds increased in dementia patients and might have a negative toxic impact on CNS functions by themselves or their degradation products (*SI Appendix*, Fig. S5) (20, 32, 39-42). Other Group B-E metabolites may be beneficial for CNS, as their quantity all declined in dementia patients. RBC-enriched Group B metabolites all containing trimethyl ammonium ion may protect CNS through their anti-oxidative and other unknown activity. Group C compounds also RBC-enriched have cellular functions implicated in energy, redox, etc. may be important for maintaining CNS brain functions. Two compounds of Group C, SAM and GSSG, had been reported to be AD- and dementia-related markers (21, 23). Group D contains 12 plasma compounds, half of which had been reported as AD-related markers. We thus speculate 7 Group A compounds pathologically enhance or lead to severe dementia such as AD. This presumed dementia deterioration by Group A factors are opposed if Group B-E metabolites is sufficiently supplied.

Correlation analysis allowed us to categorize 33 dementia markers into five subgroups (A-E) (Fig. 3). Seven Group A compounds seem to be opposed by 26 Group B-E compounds. Group A compounds were oxidized, and five of them are highly correlated, perhaps due to common relation to tryptophan degradation and/or nucleoside metabolism. Group A compounds may act as inhibitors, poisons, or wastes (20, 39). Kynurenine, quinolinic acid (40), and indoxyl-sulfate known as toxins may target to the brain and worsen dementia (32). *N*6-acetyl-lysine may be related to the accumulation of acetylated tau and alpha-tubulin in the AD brain (41, 42). Note, however, that the seven A markers increased in dementia were not found in frail subjects (16). If the change in Group A is the causal reason for dementia, then cognitive cause in frailty may be distinct from that of dementia. There seemed to be only a small overlap between dementia and frailty-related cognition markers in the outcome of patients’ metabolite analysis. Five markers are common; they are ET, *S*-methyl-ET, trimethyl-histidine, methionine, and tryptophan. Whilst aging is thought to be a major risk factor for dementia, we have not found clear age markers among dementia factors except those five.

Six Group B compounds contained trimethyl ammonium moieties, presumably acting as anti-oxidants, so that Group B may act in opposition to Group A, acting in concert to exacerbate dementia when the levels of Group B are declined. Cheah et al. (2016) indicated that the subject group with mild cognitive impairment (MCI) showed lower ET levels (17). They showed that the whole blood ET level of MCI subjects was around 0.65-fold compared to normal subjects. Kameda et al. (2020) reported that frail patients showing MCI also showed a decline of ET, S-methyl-ET, trimethyl-histidine, tryptophan, and methionine (16). In this study, the levels of ET (0.41), S-methyl-ET (0.20), and the other trimethylated ammonium compounds (0.08∼0.85) were also reduced (SI *Appendix*, Table S3). As ET is abundant, for example, in mushrooms, intervention study will be possible.

Six Group C metabolites also enriched in RBCs, include GSSG, NADP^+^, pantothenate, SAM, ATP, and gluconate, a metal (zinc) carrier. Their actions may directly or indirectly resist dementia, thus their decline would contribute to it. GSH is easily oxidized to form inactive GSSG. In this study, we only detected GSSG as GSH was difficult to stably measure. Zinc, one of the most abundant trace metals in the brain, works for dementia both positively and negatively (43, 44). Disruption of zinc homeostasis in the brain is thought to induce neurodegeneration and memory impairment (45, 46). Decreased zinc carrier gluconate may affect zinc balance in the brain. Twelve Group D compounds (amino acids, nucleosides, choline, and carnitine) are plasma enriched compounds and may underpin the actions of other metabolites for supply and degradation. Last, the two Group E compounds, caffeine and dimethyl-xanthine, also decrease in dementia. Caffeine’s beneficial effect on dementia was reported (47). Caffeine, an anti-oxidant purine, and its derivative, dimethyl-xanthine (highly correlated to each other) decline greatly in dementia subjects. Their relationship to dementia has been investigated, and caffeine may be a possible protectant against cognitive decline (28, 48), because of its anti-oxidative purine activity. Caffeine is an antagonist of adenosine (49), consistent with the present finding that adenosine belongs to Group A compounds. Caffeine and dimethyl-xanthine were correlated with trimethyl-tryptophan. Consistently, the interaction between caffeine and tryptophan in brain is known (50).

Mitochondrial dysfunction has been reported in dementia subjects (51, 52) which may explain the reduced level of ATP documented in this study in dementia subjects. ATP was clearly diminished in dementia, whereas the level was average or above in HE subjects. Our previous reports indicated that more than 98% of ATP in whole blood derives from RBC (14, 53). Therefore, a decline in ATP level in dementia subjects is thought to occur mainly in RBCs. However, it may be necessary to verify whether ATP in plasma is actually reduced. In correlation analysis, ATP concentrations were positively correlated with 10 compounds in subgroups B, C, and D (Fig. 3) so that the level of ATP may affect or be affected by concentrations of many metabolites, including oxidoreductive compounds, glutathione, betaine, ET, *S*-methyl-ET, and amino acids such as glutamine, tryptophan. Thus it is quite possible that ATP might assist anti-oxidation in the brain via many metabolites present in plasma and RBCs so that it may contribute to brain activities against dementia. NADP^+^ and GSSG may be synergistic in maintaining the level of ATP and ET (54). Hence, oxido-reductive NADP^+^, anti-oxidative glutathione, and presumably neuroprotective trimethylated ammonium compounds may all function together to sustain brain mitochondrial ATP production level against dementia. Note that glycerophosphocholine and dodecanoyl-carnitine, which also contain a trimethyl ammonium moiety, belong to Group D and may also enhance mitochondrial function.

Nine compounds possessing trimethylated ammonium ion (Table 1 and *SI Appendix*, Fig. S2) are amphipathic compounds (possessing both hydrophilic and lipophilic properties) and forming the basis of lipid polymorphism. All of them showed a sharp decline in abundance in dementia subjects. Distribution of ET in brain has been demonstrated in various mammalian species (55, 56) and human (57). The cause of its decline might be due to the rise of ROS (reactive oxygen species) in dementia patient brains. These amphipathic compounds may have similar roles, such as forming a higher-ordered, assembled structure. In addition, these compounds are abundant. They might act as major neuroprotectants or antioxidants in brain, and their levels are sensitive to both anti-oxidants and ROS. In addition, membrane defects have been observed in dementia patient brains that degrade glycerophosphocholine (58, 59). Alternatively, amphipathic moieties might deliver pertinent compounds of nM concentrations to the CNS (60). An advantage of metabolomic analysis over proteomic and genomic studies exists in the search for drugs. As these metabolites exist in the human body fluids, they may be employed for studies of drug therapy. In the present study, 33 metabolites are obvious targets for future study, the majority of which have not been studied at all relative to their clinical potential.

## Materials and Methods

### Participants

Eight dementia patients, hospitalized at the National Hospital Organization Ryukyu Hospital, in Kin-town, Okinawa, participated as subjects in this study (*SI Appendix*, Table S1). They were judged carefully by doctors at the Ryukyu Hospital to understand the study objects, contents, privacy protection, free choice of participation, and were selected. In addition, eight young and 8 elderly healthy volunteers, who live in Onna-village, Okinawa, participated as volunteers. Blood samples for LC-MS measurements were collected between 2017 and 2018.

### Diagnosis of subjects

Patients were diagnosed based on the Diagnostic and Statistical Manual of Mental Disorders, 4th edition (DSM-4) (61). Careful interviews on impairment of ADL was performed to rule out delirium or other psychiatric diseases. Blood tests and interviews for patients ruled out the possibility of the other systemic diseases. To assess cognitive ability and pathological changes, cognitive tests (HDS-R, MMSE, COGNISTAT), hippocampal atrophy by magnetic resonance imaging (MRI) for all patients, and voxel-based specific regional analysis system for Alzheimer’s disease (VSRAD) for some patients were collected (*SI Appendix*, Table S1). Both revised-Hasegawa’s dementia scale, HDS-R (62) and the mini mental state examination, MMSE (63, 64) have a maximum of 30 points, and cut-off values are 20/21 and 23/34, respectively. Dementia patients were below the cut-off values for each test, while healthy elderly people who agreed to the test were almost perfect (28∼30). The results of the Japanese version of the neurobehavioral cognitive status examination, COGNISTAT (65) and the voxel-based specific regional analysis system for Alzheimer’s disease, VSRAD (66) were done supplementally. The Z-score of VSRAD indicates the degree of atrophy of the middle temporal area, including the hippocampus (67). These clinical data were collected before January 2018 in Ryukyu Hospital. Information about patients medications is shown in Table S4.

### Ethics statement

Witten, informed consent was obtained from all participants in accordance with the Declaration of Helsinki. Consent of patient spouses or guardians were also obtained, in addition to that of the patients themselves. All experiments were performed in compliance with relevant Japanese laws and institutional guidelines. All protocols were approved by the Ethics Committee on Human Research of Ryukyu Hospital and by the Human Subjects Research Review Committee of the Okinawa Institute of Science and Technology Graduate University.

### Chemicals and reagents

Standards for metabolite identification were purchased from commercial sources, as described previously (14, 15, 53, 68). LC-MS grade acetonitrile, methanol, and ultrapure water were obtained from FUJIFILM Wako Pure Chemical Corporation (Osaka, Japan).

### Blood sample preparation

Metabolomic samples were prepared as described previously (14). Briefly, venous blood samples were collected into heparinized tubes before breakfast. Subjects were asked to ensure at least 8 h of fasting prior to sampling. During fasting, subjects took water freely. Immediately, 0.2 mL of blood were quenched in 1.8 mL of 55% methanol at -40°C. Ten nmol each of HEPES and PIPES were added to each sample to serve as standards, After brief vortexing, samples were transferred to Amicon Ultra 10-kDa cut-off filters (Millipore, Billerica, MA, USA) to remove proteins and cellular debris. After sample concentration by vacuum evaporation, each sample was re-suspended in 40 µL of 50% acetonitrile, and 1 µL was used for each injection into the LC-MS system, as described.

### LC-MS analysis

Non-targeted LC-MS conditions were as described previously (14, 16). Briefly, LC-MS data were obtained using an Ultimate 3000 DGP-3600RS HPLC system (Thermo Fisher Scientific, Waltham, MA, USA) coupled to an LTQ Orbitrap mass spectrometer (Thermo Fisher Scientific, Waltham, MA, USA). LC separation was performed on a ZIC-pHILIC column (Merck SeQuant, Umea, Sweden; 150 mm × 2.1 mm, 5 µm particle size). Acetonitrile (A) and 10 mM ammonium carbonate buffer, pH 9.3 (B) were used as the mobile phase, with a linear gradient from 80-20% A over 30 min, at a flow rate of 100 µL/mL. The mass spectrometer was operated in full-scan mode with a 100-1000 m/z scan rate and automatic data-dependent MS/MS fragmentation scans.

### LC-MS data processing and peak characteristics

Peak areas of metabolites of interest were measured using MZmine 2 software (69). Data analytical procedures and parameters have been described previously (15). We analyzed 124 blood metabolites that were confirmed using standards or MS/MS analysis (Dataset S1). According to their peak areas, metabolite abundances were classified into 3 groups (H, M, and L). H denotes compounds with high peak areas (>10^8^ AU), M with medium peak areas (10^7^∼10^8^ AU) and L with low peak areas (<10^7^ AU) (*SI Appendix*, Table S2).

### Statistical analysis

Peak data processed with MZmine 2 were exported into spreadsheet format and analyzed with R statistical software (http://www.r-project.org). Statistical analysis was performed using the Mann Whitney U-test. Statistical significance was established at *P* < 0.05. *Q*-values were calculated using the Benjamini-Hochberg method. The plot of principal components (PC) was generated with SIMCA-P+ software (Umetrics Inc., Umea, Sweden). The heatmap represents standardized abundance data for each metabolite. T-scores were calculated from the following formula: T-score = [(sample peak area – average of population peak area) × 10/standard deviation of population peak area] + 50. Therefore, the mean and standard deviation are 50 and 10, respectively.

## Data availability

Raw LC-MS data in mzML format are accessible via the MetaboLights repository (http://www.ebi.ac.uk/metabolights). Data for the 24 subjects are available under accession number MTBLS2109.

## Acknowledgments

We are greatly indebted to Ryukyu Hospital and volunteers of Onna-village, without which the study would not have been possible. We are grateful to Ms. Hiromi Karimata for excellent assistance in coordinating clinical data and documents. We thank Ms. Lisa Uehara, Dr. Michiko Suma, Ms. Ayaka Mori, and Ms. Yuria Tahara for providing excellent technical assistance, Ms. Kaori Serakaki for illustration (Fig. S5), Dr. Reiko Sugiura for discussions on clinical data, and Dr. Steven D. Aird for editing the manuscript. We are greatly indebted to generous support from the Okinawa Institute of Science and Technology Graduate University and its POC program fund.

## Notes

### Competing Interest Statement

The authors have declared no competing interest.

