## Supplementary material for "Whole blood metabolomics of dementia patients reveal classes of disease-linked metabolites": Figures S1-S5 and Tables S1-S4

**This PDF file includes:**

Figures S1 to S5
Tables S1 to S4
Legend for Dataset S1

**Other supplementary materials for this manuscript include the following:**

Dataset S1

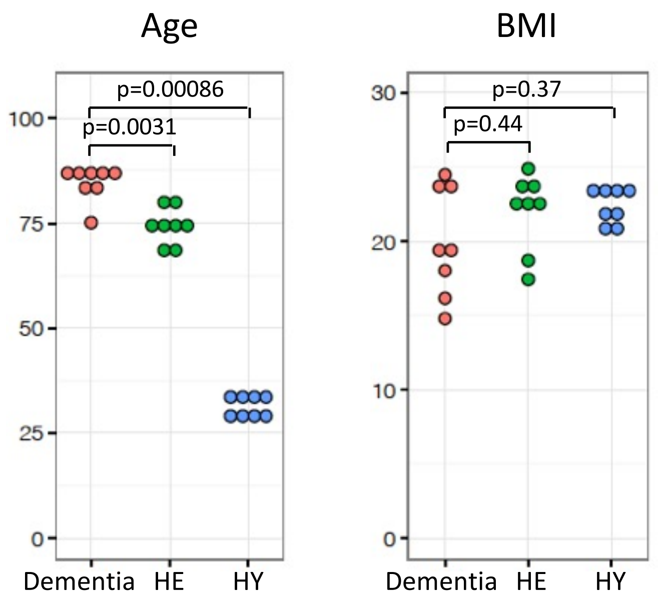


**Fig. S1.** Patients with dementia, healthy elderly (HE), and healthy young (HY) volunteers. Age and BMI (body mass index) are schematically shown. Ages of dementia patients ranged from 75 to 88 years and their BMIs varied from 14.8 to 24.5. The 8 HE subjects ranged in age from 67-80 with BMIs ranging from 17-25. The 8 HY subjects were between 28 and 34 years with BMIs 21 to 24.

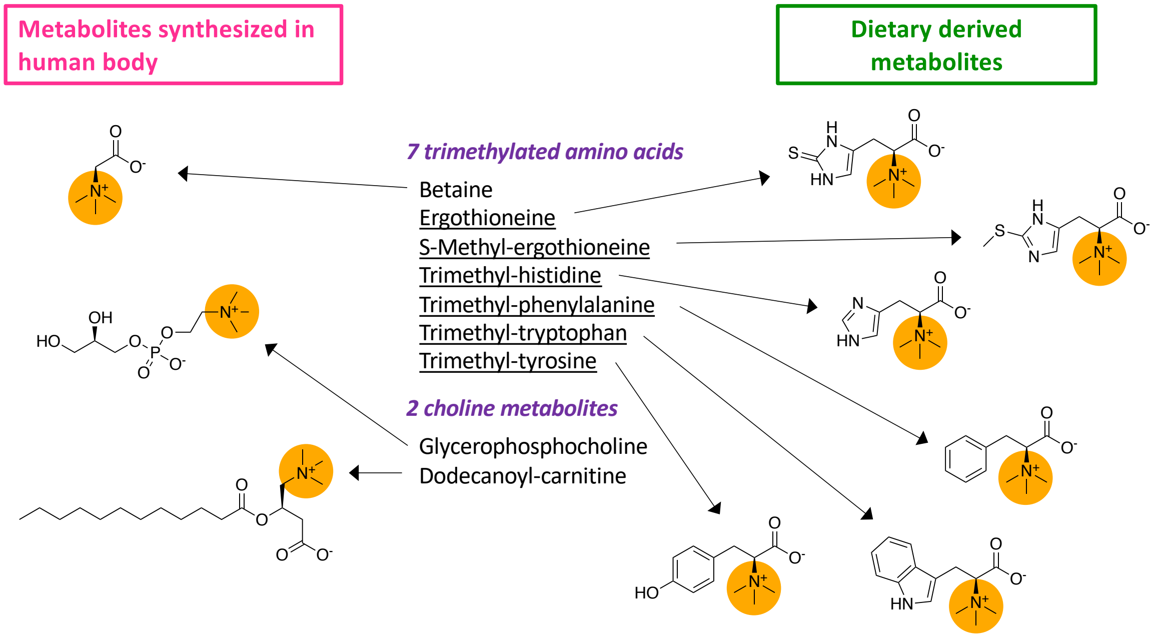


**Fig. S2.** In dementia patients, the abundance of various trimethylated compounds decreased in blood. Three trimethylated compounds (betaine, glycerophosphocholine, dodecanoyl-carnitine) are synthesized in the human body, whereas six other compounds (ergothioneine, *S*-methyl-ergothioneine, trimethyl-histidine, trimethyl-phenylalanine, trimethyl-tryptophan, trimethyl-tyrosine) are derived from food. These nine compounds decreased in whole blood of dementia patients.

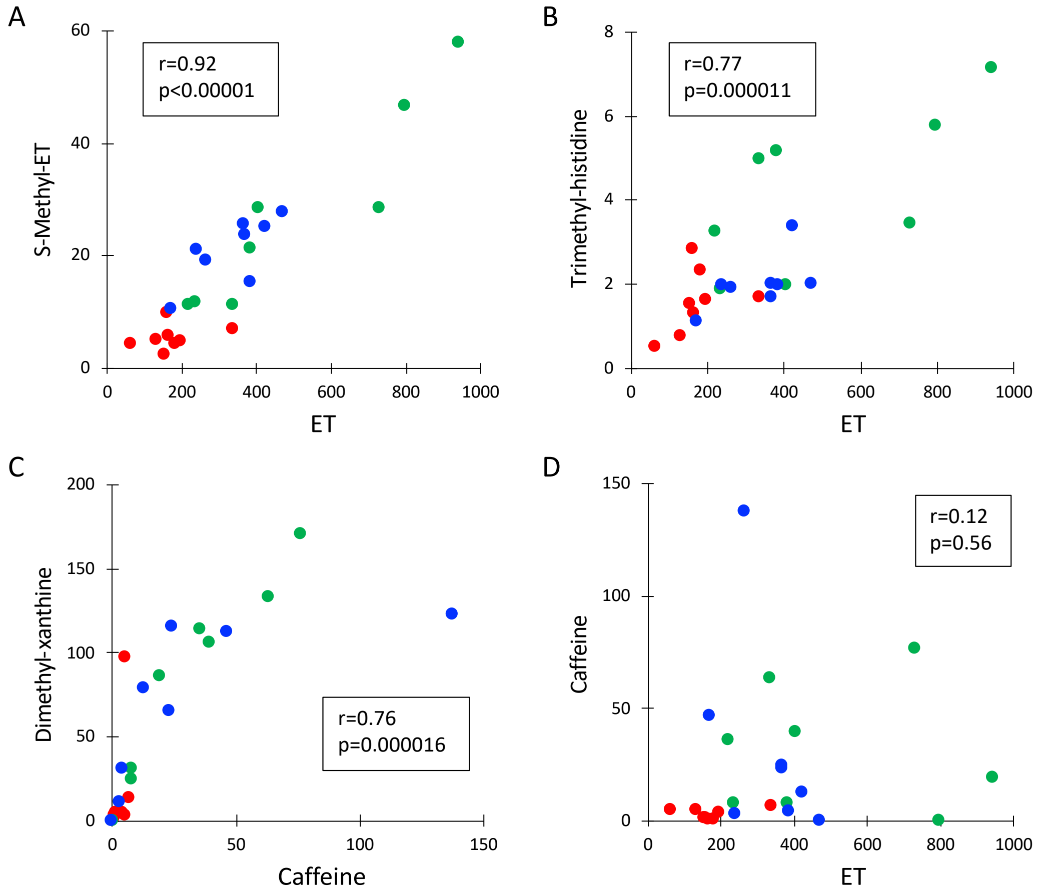


**Fig. S3.** Scatter plots showing high correlations between related metabolites. Metabolites related to ergothioneine biosynthesis (*A* and *B*) and caffeine catabolism (*C*) are displayed. Note, in contrast, that metabolically unrelated compounds, ET and caffeine, are not correlated at all (*D*). Metabolite abundance (peak area, $x10^6$ AU) of dementia patients (red dots), HE (green), and HY (blue) are represented. Pearson's correlation coefficient r and p-value are shown in boxes.

| 33 dementia markers | | | HY | | | | | | | |
|---|---|---|---|---|---|---|---|---|---|---|
| | | | #17 | #18 | #19 | #20 | #21 | #22 | #23 | #24 |
| Higher in dementia (7) | A | Quinolinic acid | 40.3 | 40.5 | 49.8 | 50.1 | 50.0 | 42.1 | 54.5 | 45.4 |
| | | Dimethyl-guanosine | 41.1 | 32.9 | 40.4 | 52.4 | 53.8 | 40.3 | 44.2 | 45.3 |
| | | Pseudouridine | 45.3 | 38.0 | 35.8 | 45.0 | 43.3 | 46.1 | 43.6 | 47.2 |
| | | Indoxyl-sulfate | 36.6 | 47.8 | 43.1 | 38.8 | 55.9 | 45.7 | 52.6 | 48.7 |
| | | Kynurenine | 44.3 | 29.5 | 50.3 | 52.1 | 61.1 | 48.0 | 53.5 | 45.1 |
| | | N6-Acetyl-lysine | 61.0 | 53.5 | 46.1 | 46.9 | 48.7 | 43.7 | 43.7 | 43.4 |
| | | Adenosine | 48.6 | 48.0 | 52.9 | 59.3 | 86.2 | 48.6 | 48.2 | 49.5 |
| Lower in dementia (26) | B | S-Methyl-ergothioneine | 55.3 | 54.1 | 51.0 | 57.2 | 48.2 | 55.7 | 44.6 | 52.3 |
| | | Ergothioneine | 53.9 | 51.4 | 46.6 | 56.1 | 52.2 | 51.3 | 42.3 | 45.4 |
| | | Trimethyl-histidine | 54.7 | 44.5 | 45.9 | 46.4 | 46.3 | 46.5 | 41.2 | 46.3 |
| | | Trimethyl-tryptophan | 58.5 | 42.8 | 62.5 | 45.7 | 50.3 | 43.0 | 42.7 | 43.8 |
| | | Trimethyl-phenylalanine | 47.0 | 47.0 | 47.1 | 47.0 | 46.9 | 46.9 | 46.9 | 47.1 |
| | | Trimethyl-tyrosine | 46.4 | 53.8 | 50.7 | 57.0 | 53.5 | 48.5 | 45.5 | 49.4 |
| | C | Pantothenate | 44.7 | 41.2 | 55.2 | 53.8 | 52.3 | 45.4 | 64.5 | 41.0 |
| | | Gluconate | 47.2 | 36.8 | 48.6 | 59.8 | 48.4 | 45.5 | 48.1 | 55.6 |
| | | S-Adenosyl-methionine | 84.3 | 66.5 | 52.1 | 51.9 | 44.0 | 46.1 | 54.2 | 59.9 |
| | | NADP+ | 64.4 | 50.5 | 61.3 | 73.7 | 59.1 | 54.2 | 50.6 | 55.1 |
| | | Glutathione disulfide | 58.2 | 49.8 | 48.6 | 65.7 | 47.1 | 43.8 | 47.5 | 61.0 |
| | | ATP | 49.3 | 46.8 | 42.9 | 67.6 | 63.2 | 52.1 | 35.3 | 64.6 |
| | D | Methionine | 41.6 | 39.4 | 55.9 | 62.4 | 63.4 | 47.1 | 52.4 | 57.5 |
| | | Tryptophan | 42.4 | 41.7 | 59.6 | 64.8 | 63.0 | 45.5 | 52.4 | 58.6 |
| | | Glutamine | 44.1 | 47.6 | 37.4 | 67.6 | 55.3 | 37.9 | 49.1 | 56.1 |
| | | Betaine | 43.7 | 38.7 | 42.2 | 62.9 | 72.9 | 55.5 | 36.9 | 53.7 |
| | | Phenylalanine | 36.3 | 37.4 | 46.6 | 54.6 | 69.5 | 40.4 | 46.6 | 55.4 |
| | | Tyrosine | 37.7 | 39.1 | 69.1 | 67.4 | 57.6 | 39.0 | 52.1 | 49.5 |
| | | Histidine | 44.7 | 44.7 | 42.2 | 53.4 | 53.3 | 49.2 | 46.1 | 55.8 |
| | | Uridine | 54.0 | 43.2 | 35.5 | 52.9 | 53.4 | 56.8 | 54.1 | 71.5 |
| | | Keto(iso)leucine | 44.3 | 39.1 | 37.0 | 66.4 | 43.5 | 53.1 | 49.9 | 68.1 |
| | | Glycerophosphocholine | 57.0 | 53.1 | 41.5 | 74.1 | 81.0 | 43.8 | 48.1 | 49.7 |
| | | 2-Hydroxybutyrate | 55.7 | 34.7 | 39.6 | 42.2 | 44.2 | 39.6 | 49.6 | 63.6 |
| | | Dodecanoyl-carnitine | 67.9 | 47.7 | 48.4 | 42.8 | 40.6 | 40.8 | 68.3 | 68.2 |
| | E | Caffeine | 47.1 | 50.8 | 85.6 | 43.3 | 44.6 | 50.4 | 57.6 | 44.2 |
| | | Dimethyl-xanthine | 54.3 | 60.9 | 62.3 | 39.8 | 45.4 | 51.8 | 60.5 | 41.8 |

**Fig. S4.** A heatmap of standardized metabolite abundances of 33 dementia markers from healthy young (HY) subjects.

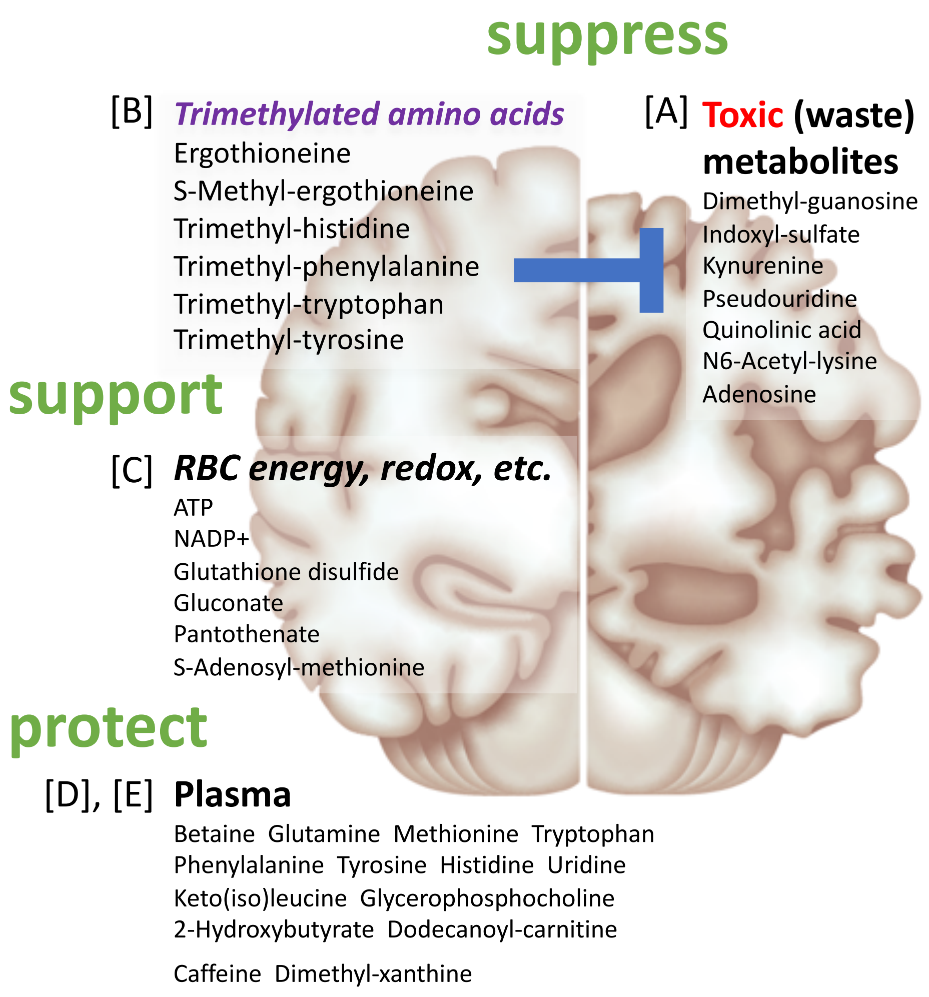


**Fig. S5.** Subclass A compounds may inhibit normal brain functions, leading to dementia, whereas subclasses B-E oppose dementia, possibly protecting the brain. Some compounds of group D enriched in plasma were previously shown to be dementia markers. Compounds of groups B and C enriched in RBC are mostly novel dementia markers (see text for explanation).

**Table S1.** Characteristics for dementia patients and healthy young (HY) and elderly (HE) subjects.

| Group | Subject | Age | Gender | BMI | Cognitive test | | | Brain imaging | |
|---|---|---|---|---|---|---|---|---|---|
| | | | | | HDS-R score | MMSE score | COGNISTAT total score | MRI finding | VSRAD z-score |
| Dementia | #1 | 87 | F | 16.2 | 13 | 11 | 45 | HA, CA | - |
| | #2 | 85 | F | 19.2 | 10 | 9 | - | HA, WML | 4.05 |
| | #3 | 82 | M | 19.7 | 6 | 12 | - | HA, CA | - |
| | #4 | 87 | M | 14.8 | 12 | 10 | - | HA, WML, OBA | 6.27 |
| | #5 | 87 | F | 18.0 | 5 | 7 | - | HA, CA, WML | 6.31 |
| | #6 | 86 | F | 24.5 | 18 | 22 | 88 | HA, CA, OLI | 3.55 |
| | #7 | 88 | M | 23.4 | 17 | 21 | 63 | HA, CA, WHL, OLI | 2.03 |
| | #8 | 75 | M | 24.0 | 11 | - | - | HA, OBA, OLI | - |
| HE | #9 | 67 | M | 22.3 | 30 | 29 | - | - | - |
| | #10 | 74 | M | 23.6 | - | - | - | - | - |
| | #11 | 73 | M | 23.9 | - | - | - | - | - |
| | #12 | 70 | F | 22.1 | 28 | 29 | - | - | - |
| | #13 | 75 | F | 24.9 | 30 | 30 | - | - | - |
| | #14 | 76 | M | 23.0 | 28 | 29 | - | - | - |
| | #15 | 80 | F | 17.4 | 30 | 30 | - | - | - |
| | #16 | 80 | F | 18.7 | 28 | 30 | - | - | - |
| HY | #17 | 30 | M | 23.1 | - | - | - | - | - |
| | #18 | 28 | F | 21.0 | - | - | - | - | - |
| | #19 | 33 | F | 21.9 | - | - | - | - | - |
| | #20 | 34 | M | 21.9 | - | - | - | - | - |
| | #21 | 34 | F | 20.7 | - | - | - | - | - |
| | #22 | 33 | M | 23.0 | - | - | - | - | - |
| | #23 | 30 | F | 23.3 | - | - | - | - | - |
| | #24 | 30 | M | 23.9 | - | - | - | - | - |

Dementia was diagnosed by DSM-4 criteria, cognitive tests, and brain imaging (details in the Methods section). BMI, body mass index; HDS-R, Hasegawa's dementia scale-revised; MMSE, mini mental state examination; MRI, magnetic resonance imaging; VSRAD, voxel-based specific regional analysis system for Alzheimer's disease; COGNISTAT, neurobehavioural cognitive status examination; HA, hippocampal atrophy; CA, cerebral atrophy; WML, white matter lesions; OBA, overall brain atrophy; OLI, old lacunar infarction. Short line (-) indicates *not examined*.

**Table S2.** List of 124 identified blood metabolites

| ***Category***/Compound | | Peak abundance | CV | | Dementia/HE | | |
|---|---|---|---|---|---|---|---|
| | | | present study | Chaleckis *et al.* (14) | *P*-value | Ratio | *Q*-value |
| ***Nucleotides (9)*** | | | | | | | |
| * | ATP | H | 0.20 | 0.17 | 0.0047 | 0.83 | 0.0171 |
| | ADP | H-M | 0.30 | 0.27 | 0.1949 | 0.83 | |
| | AMP | M | 0.35 | 0.33 | 0.1605 | 0.80 | |
| | GTP | M | 0.21 | 0.32 | 0.5737 | 1.10 | |
| | UMP | M | 0.20 | 0.88 | 0.1304 | 1.24 | |
| | GDP | M-L | 0.26 | 0.38 | 0.4418 | 1.21 | |
| | CTP | L | 0.38 | 0.33 | 0.2345 | 1.22 | |
| | UDP | L | 0.27 | 0.31 | 0.1304 | 1.28 | |
| | UTP | L | 0.32 | 0.44 | 0.0650 | 1.38 | |
| ***Nucleosides, nucleobases, and derivatives (12)*** | | | | | | | |
| * | Caffeine | H-L | 1.49 | 0.92 | 0.0104 | 0.09 | 0.0344 |
| * | Dimethyl-xanthine | H-L | 0.98 | 0.61 | 0.0281 | 0.04 | 0.0422 |
| | Urate | M | 0.21 | 0.28 | 0.0650 | 0.85 | |
| | Adenine | M-L | 0.43 | 0.53 | 0.2786 | 0.77 | |
| | Cytidine | M-L | 0.34 | 0.33 | 0.0650 | 0.67 | |
| | Hypoxanthine | M-L | 0.46 | 0.35 | 0.8785 | 0.93 | |
| | *N*-Methyl-adenosine | M-L | 0.31 | 0.29 | 0.7985 | 0.92 | |
| * | Pseudouridine | M-L | 0.31 | - | 0.0148 | 1.19 | 0.0304 |
| | Xanthine | M-L | 1.93 | 0.56 | 0.9591 | 0.91 | |
| * | Adenosine | L | 0.47 | 0.49 | 0.0379 | 1.51 | 0.0481 |
| * | Dimethyl-guanosine | L | 0.40 | 0.46 | 0.0499 | 1.45 | 0.0549 |
| * | Uridine | L | 0.30 | 0.34 | 0.0030 | 0.72 | 0.0122 |
| ***Vitamins and coenzymes (5)*** | | | | | | | |
| | Nicotinamide | H-M | 0.30 | 0.56 | 0.9591 | 1.04 | |
| | NAD+ | M | 0.35 | 0.30 | 0.4418 | 1.13 | |
| | 4-Aminobenzoate | M-L | 3.47 | 2.18 | 0.5054 | 0.18 | |
| * | Pantothenate | M-L | 0.46 | 0.82 | 0.0148 | 0.57 | 0.0287 |
| * | NADP+ | L | 0.33 | 0.36 | 0.0104 | 0.70 | 0.0312 |
| ***Nucleotide-sugar derivatives (4)*** | | | | | | | |
| | UDP-N-acetyl-glucosamine | M | 0.55 | 0.64 | 0.5737 | 0.85 | |
| | UDP-glucose | M | 0.24 | 0.24 | 0.5737 | 0.95 | |
| | GDP-glucose | M-L | 0.37 | 0.53 | 0.4418 | 0.87 | |
| | UDP-glucuronate | L | 0.38 | 0.63 | 0.9591 | 0.86 | |
| ***Sugar phosphates (11)*** | | | | | | | |
| | Diphosphoglycerate | H | 0.18 | 0.24 | 0.8785 | 0.96 | |
| | Fructose-1,6-diphosphate | H-M | 0.19 | - | 0.2345 | 1.15 | |
| | Glucose-6-phosphate | M | 0.19 | 0.29 | 0.4418 | 1.11 | |
| | Glyceraldehyde-3-phosphate | M | 0.38 | 0.99 | 0.0650 | 0.59 | |
| | Fructose-6-phosphate | M-L | 0.21 | 0.24 | 0.5054 | 1.23 | |
| | Glycerol-phosphate | M-L | 0.28 | 0.31 | 0.6454 | 0.99 | |
| | Phosphoglycerate | M-L | 0.22 | 0.29 | 0.5054 | 0.93 | |
| | 6-Phosphogluconate | L | 0.20 | 0.30 | 0.9591 | 0.96 | |
| | Pentose-phosphate | L | 0.22 | 0.34 | 0.0650 | 1.21 | |
| | Phosphoenolpyruvate | L | 0.18 | - | 0.2345 | 0.88 | |
| | Sedoheptulose-7-phosphate | L | 0.24 | 0.52 | 0.0830 | 1.25 | |
| ***Sugar derivatives (4)*** | | | | | | | |

| | | | | | | | |
|---|---|---|---|---|---|---|---|
| | 1,5-Anhydroglucitol | M | 0.30 | 0.46 | 0.6454 | 1.10 | |
| * | Gluconate | M | 0.17 | 0.33 | 0.0379 | 0.85 | 0.0464 |
| | *N*-Acetyl-glucosamine | M | 0.25 | 0.26 | 0.6454 | 0.88 | |
| | myo-Inositol | M-L | 0.32 | 0.24 | 0.3823 | 0.86 | |
| ***Choline and ethanolamine derivatives (7)*** | | | | | | | |
| | Choline | H-M | 0.74 | - | 0.1949 | 0.64 | |
| * | Glycerophosphocholine | H-M | 0.49 | 0.47 | 0.0148 | 0.66 | 0.0271 |
| | Phosphocholine | H-M | 0.28 | - | 0.7209 | 1.06 | |
| | CDP-choline | M-L | 0.38 | 0.41 | 0.1605 | 0.80 | |
| | CDP-ethanolamine | L | 0.42 | 0.43 | 0.6454 | 0.88 | |
| | Glycerophosphoethanolamine | L | 0.35 | - | 0.1949 | 0.83 | |
| | Phosphoethanolamine | L | 0.27 | - | 0.4418 | 1.04 | |
| ***Organic acids (10)*** | | | | | | | |
| * | 2-Hydroxybutyrate | M-L | 0.34 | - | 0.0499 | 0.79 | 0.0531 |
| | Chenodeoxycholate | M-L | 1.67 | 1.33 | 0.1949 | 1.99 | |
| | Citrate | M-L | 0.18 | 0.31 | 0.9591 | 1.03 | |
| | Glycochenodeoxycholate | M-L | 2.44 | 1.20 | 0.5737 | 1.59 | |
| | 2-Oxoglutarate | L | 0.46 | 0.54 | 0.1304 | 0.73 | |
| | 3-Hydroxybutyrate | L | 0.95 | - | 0.3282 | 0.68 | |
| | Citramalate | L | 0.24 | 0.36 | 0.1605 | 0.87 | |
| | Glycerate | L | 0.37 | 0.43 | 0.5737 | 0.92 | |
| | Malate | L | 0.20 | 0.20 | 0.2345 | 0.91 | |
| | Succinate | L | 0.36 | 0.60 | 0.3823 | 0.81 | |
| ***Antioxidants (2)*** | | | | | | | |
| * | Glutathione disulfide | H | 0.14 | 0.18 | 0.0104 | 0.82 | 0.0286 |
| * | Ergothioneine | H-M | 0.64 | 0.63 | 0.0011 | 0.41 | 0.0090 |
| ***Standard amino acids (17)*** | | | | | | | |
| | Arginine | H | 0.14 | 0.29 | 0.7985 | 0.96 | |
| * | Glutamine | H | 0.14 | 0.20 | 0.0207 | 0.84 | 0.0341 |
| * | Phenylalanine | H | 0.18 | 0.17 | 0.0281 | 0.73 | 0.0404 |
| | Proline | H | 0.28 | 0.42 | 0.1304 | 0.86 | |
| | Glutamate | H-M | 0.16 | 0.28 | 0.1304 | 0.84 | |
| | Lysine | H-M | 0.18 | 0.39 | 0.8785 | 0.96 | |
| * | Methionine | H-M | 0.16 | 0.28 | 0.0379 | 0.80 | 0.0447 |
| | Threonine | H-M | 0.19 | 0.42 | 0.5054 | 0.96 | |
| * | Tryptophan | H-M | 0.22 | 0.24 | 0.0281 | 0.80 | 0.0387 |
| * | Tyrosine | H-M | 0.21 | 0.27 | 0.0104 | 0.75 | 0.0264 |
| | Asparagine | M | 0.20 | 0.23 | 0.1304 | 0.90 | |
| * | Histidine | M | 0.24 | 0.19 | 0.0379 | 0.80 | 0.0432 |
| | Isoleucine | M | 0.19 | 0.32 | 0.4418 | 0.86 | |
| | Leucine | M | 0.18 | 0.31 | 0.0830 | 0.83 | |
| | Serine | M | 0.20 | 0.33 | 0.1304 | 0.84 | |
| | Valine | M | 0.20 | 0.48 | 0.0650 | 0.87 | |
| | Aspartate | M-L | 0.37 | 0.50 | 1.0000 | 0.91 | |
| ***Methylated amino acids (14)*** | | | | | | | |
| * | Betaine | H | 0.16 | 0.51 | 0.0019 | 0.85 | 0.0123 |
| | Butyro-betaine | H-M | 0.23 | 0.35 | 0.5054 | 0.92 | |
| | Dimethyl-proline | H-M | 0.53 | 0.79 | 0.8785 | 0.95 | |
| | Trimethyl-lysine | H-M | 0.30 | 0.38 | 0.5737 | 1.04 | |
| * | Trimethyl-tryptophan | H-L | 1.09 | 1.67 | 0.0003 | 0.10 | 0.0051 |

| | | | | | | |
|---|---|---|---|---|---|---|
| Dimethyl-arginine | M | 0.24 | 0.31 | 0.7209 | 1.11 | |
| *N*1-Methyl-histidine | M | 0.40 | 0.30 | 1.0000 | 0.92 | |
| Dimethyl lysine | M-L | 0.64 | 0.44 | 0.0830 | 1.58 | |
| *N*3-Methyl-histidine | M-L | 0.49 | 0.30 | 0.9591 | 1.01 | |
| *N*6-Methyl-lysine | M-L | 0.71 | 0.73 | 0.7209 | 0.84 | |
| * *S*-Methyl-ergothioneine | M-L | 0.77 | 0.63 | 0.0002 | 0.20 | 0.0053 |
| * Trimethyl-phenylalanine | M-L | 3.17 | 1.18 | 0.0006 | 0.23 | 0.0068 |
| * Trimethyl-tyrosine | M-L | 1.09 | 2.50 | 0.0019 | 0.08 | 0.0102 |
| * Trimethyl-histidine | L | 0.64 | 0.57 | 0.0019 | 0.37 | 0.0088 |
| ***Acetylated amino acids (4)*** | | | | | | |
| *N*-Acetyl-arginine | L | 0.52 | 0.62 | 0.7985 | 1.08 | |
| *N*-Acetyl-aspartate | L | 0.23 | 0.58 | 0.2345 | 1.09 | |
| *N*2-Acetyl-lysine | L | 0.30 | 0.50 | 0.6454 | 0.86 | |
| * *N*6-Acetyl-lysine | L | 0.79 | 0.43 | 0.0207 | 1.47 | 0.0325 |
| ***Other amino acids (16)*** | | | | | | |
| Creatine | M | 0.33 | 0.29 | 0.0650 | 1.39 | |
| Taurine | M | 0.18 | 0.37 | 1.0000 | 1.05 | |
| Creatinine | H | 0.16 | 0.35 | 0.4418 | 0.93 | |
| * Indoxyl-sulfate | M | 0.46 | 0.59 | 0.0148 | 1.93 | 0.0256 |
| Hippurate | M-L | 0.83 | 1.01 | 0.1605 | 0.47 | |
| * Keto(iso)leucine | M-L | 0.26 | - | 0.0104 | 0.77 | 0.0245 |
| * Kynurenine | M-L | 0.16 | 0.48 | 0.0281 | 1.12 | 0.0371 |
| Ophthalmic acid | M-L | 0.41 | 0.43 | 0.1949 | 0.71 | |
| Ornithine | M-L | 0.29 | 0.48 | 0.3823 | 1.17 | |
| 4-Guanidinobutanoate | L | 0.71 | 2.05 | 0.2786 | 0.60 | |
| Acetyl-carnosine | L | 0.65 | 1.07 | 0.2786 | 0.60 | |
| Citrulline | L | 0.27 | 0.30 | 0.4418 | 0.95 | |
| Phosphocreatine | L | 0.47 | 0.48 | 0.7985 | 1.18 | |
| * Quinolinic acid | L | 0.48 | 1.18 | 0.0499 | 1.79 | 0.0514 |
| *S*-Adenosyl-homocysteine | L | 0.34 | 0.83 | 0.1949 | 1.11 | |
| * *S*-Adenosyl-methionine | L | 1.02 | 0.88 | 0.0104 | 0.35 | 0.0229 |
| ***Carnitines (9)*** | | | | | | |
| Acetyl-carnitine | H | 0.28 | 0.41 | 0.5737 | 0.96 | |
| Carnitine | H | 0.14 | 0.20 | 0.2786 | 1.10 | |
| Propionyl-carnitine | H-M | 0.33 | 0.41 | 0.5737 | 0.87 | |
| (iso)Butyryl-carnitine | M-L | 0.45 | 0.37 | 0.8785 | 1.12 | |
| Decanoyl-carnitine | M-L | 0.75 | 1.11 | 0.2345 | 0.67 | |
| Hexanoyl-carnitine | M-L | 0.50 | 0.76 | 0.3282 | 0.73 | |
| Octanoyl-carnitine | M-L | 0.78 | 1.02 | 0.5737 | 0.69 | |
| (iso)Valeryl-carnitine | M-L | 0.35 | 1.98 | 0.1605 | 0.81 | |
| * Dodecanoyl-carnitine | L | 0.91 | 1.11 | 0.0499 | 0.15 | 0.0499 |

*P*-values between patients with dementia and HE were calculated using a Mann-Whitney U-test. Asterisks indicate 33 compounds that showed significant differences ($P < 0.05$) between dementia patients and HE. Coefficients of variation (CV) from the present and a previous study (ref 14) are listed. They are mostly in agreement. Short line (-) indicates not reported. The peak ratio was calculated using the median of peak abundance in dementia patients and HE, respectively. Fifty-one RBC-enriched compounds are underlined (ref 14). To estimate the false discovery rate, *Q*-values were also calculated. *Q*-values were mostly consistent with *P*-values of 33 metabolites.

**Table S3.** The ratio of the metabolite levels in whole blood between healthy controls, and patients with mild cognitive impairment (MCI), frailty, and dementia.

| | MCI<br>Cheah et al. (ref 17) | Frailty<br>Kameda et al. (ref 16) | Dementia<br>present study |
|---|---|---|---|
| Ergothioneine | 0.65 | 0.53 | 0.41 |
| S-Methyl-ergothioneine | not reported | 0.48 | 0.20 |
| Trimethyl-histidine | | 0.53 | 0.37 |
| Tryptophan | | 0.80 | 0.80 |
| Methionine | | 0.81 | 0.80 |

The ratio was calculated by dividing the quantitative value of a patient by that of healthy controls.

**Table S4.** Medication information about dementia patients.

| Subject | Medication |
|---|---|
| #1 | Mem, Ari, Esz, Ram, Tul, Bromhexine hydrochloride, Theophylline, Montelukast sodium, Salmeterol xinafoate, Celecoxib |
| #2 | Mem, Ari, Esz, Ram, Val, Nif, Rab, Limaprost alfadex |
| #3 | Mem, Esz, Val, Nif, Sodium ferrous citrate, Magnesium oxide |
| #4 | Mem, Suv, Yokukansan, Potassium gluconate |
| #5 | Galantamine hydrobromide, Esz, Ram |
| #6 | Mem, Esz, Suv |
| #7 | Ram, Tul, Nicorandil, Isosorbide dinitrate, Olmesartan medoxomil, Febuxostat, Tamsulosin hydrochloride, Imidafenacin, Goshajinkigan, Lubiprostone |
| #8 | Rab, Amlodipine besilate |

Mem, Memantine hydrochloride; Ari, Aripiprazole; Esz, Eszopiclone; Ram, Ramelteon; Tul, Tulobuterol; Val, Valsartan; Nif, Nifedipine; Rab, Rabeprazole sodium; Suv, Suvorexant.

**Dataset S1 (separate file).** Compounds were identified using either commercially available standards (STD) or by analysis of MS/MS spectra (MS/MS), if no standard was available.
