## Supplementary material for "Whole blood metabolomics of dementia patients reveal classes of disease-linked metabolites": Dataset S1

**Dataset S1.** List of 124 identified compounds in blood samples

| Compounds | Status | Formula | Ionization | Theoretical m/z | Detected m/z | error (ppm) | RT (min) |
|---|---|---|---|---|---|---|---|
| 1,5-Anhydroglucitol | STD | C6H12O5 | [M-H]- | 163.0612 | 163.0612 | -0.1 | 9.4 |
| 2-Hydroxybutyrate | STD | C4H8O3 | [M-H]- | 103.0401 | 103.0404 | 3.2 | 4.4 |
| 2-Oxoglutarate | STD | C5H6O5 | [M-H]- | 145.0142 | 145.0141 | -0.7 | 14.3 |
| 3-Hydroxybutyrate | STD | C4H8O3 | [M-H]- | 103.0401 | 103.0404 | 3.1 | 5.6 |
| 4-Aminobenzoate | STD | C7H7NO2 | [M+H]+ | 138.0555 | 138.0545 | -7.5 | 7.0 |
| 4-Guanidinobutanoate | STD | C5H11N3O2 | [M+H]+ | 146.0930 | 146.0921 | -6.0 | 14.4 |
| 6-Phosphogluconate | STD | C6H13O10P | [M-H]- | 275.0174 | 275.0166 | -2.8 | 17.6 |
| Acetyl-carnitine | STD | C9H17NO4 | [M+H]+ | 204.1236 | 204.1227 | -4.4 | 8.6 |
| Acetyl-carnosine | STD | C11H16N4O4 | [M+H]+ | 269.1250 | 269.1240 | -3.7 | 6.0 |
| Adenine | STD | C5H5N5 | [M+H]+ | 136.0623 | 136.0612 | -7.9 | 6.5 |
| Adenosine | STD | C10H13N5O4 | [M+H]+ | 268.1046 | 268.1035 | -4.1 | 6.1 |
| ADP | STD | C10H15N5O10P2 | [M+H]+ | 428.0372 | 428.0359 | -3.1 | 15.0 |
| AMP | STD | C10H14N5O7P | [M+H]+ | 348.0709 | 348.0698 | -3.3 | 12.9 |
| Arginine | STD | C6H14N4O2 | [M+H]+ | 175.1195 | 175.1186 | -5.4 | 26.7 |
| Asparagine | STD | C4H8N2O3 | [M+H]+ | 133.0613 | 133.0603 | -7.7 | 14.2 |
| Aspartate | STD | C4H7NO4 | [M-H]- | 132.0302 | 132.0302 | -0.5 | 13.9 |
| ATP | STD | C10H16N5O13P3 | [M+H]+ | 508.0036 | 508.0018 | -3.4 | 16.3 |
| Betaine | STD | C5H11NO2 | [M+H]+ | 118.0868 | 118.0858 | -8.8 | 8.7 |
| Butyro-betaine | STD | C7H15NO2 | [M+H]+ | 146.1181 | 146.1171 | -6.9 | 12.3 |
| (iso)Butyryl-carnitine | STD | C11H21NO4 | [M+H]+ | 232.1549 | 232.1540 | -3.8 | 5.9 |
| Caffeine | STD | C8H10N4O2 | [M+H]+ | 195.0882 | 195.0873 | -4.6 | 3.7 |
| Carnitine | STD | C7H15NO3 | [M+H]+ | 162.1130 | 162.1120 | -6.0 | 12.0 |
| CDP-choline | STD | C14H26N4O11P2 | [M+H]+ | 489.1152 | 489.1134 | -3.5 | 14.9 |
| CDP-ethanolamine | STD | C11H20N4O11P2 | [M+H]+ | 447.0682 | 447.0668 | -3.0 | 16.0 |
| Chenodeoxycholate | STD | C24H40O4 | [M-H]- | 391.2854 | 391.2849 | -1.1 | 3.1 |
| Choline | STD | C5H13NO | [M+H]+ | 104.1075 | 104.1065 | -10.1 | 19.5 |
| Citramalate | STD | C5H8O5 | [M-H]- | 147.0299 | 147.0297 | -1.1 | 14.1 |
| Citrate | STD | C6H8O7 | [M-H]- | 191.0197 | 191.0194 | -1.8 | 17.8 |
| Citrulline | STD | C6H13N3O3 | [M-H]- | 174.0884 | 174.0883 | -0.5 | 15.3 |
| Creatine | STD | C4H9N3O2 | [M-H]- | 130.0622 | 130.0622 | -0.1 | 13.8 |
| Creatinine | STD | C4H7N3O | [M+H]+ | 114.0667 | 114.0657 | -9.2 | 6.7 |
| CTP | STD | C9H16N3O14P3 | [M-H]- | 481.9772 | 481.9771 | -0.3 | 18.0 |
| Cytidine | STD | C9H13N3O5 | [M+H]+ | 244.0933 | 244.0925 | -3.3 | 9.7 |
| Decanoyl-carnitine | STD | C17H33NO4 | [M+H]+ | 316.2488 | 316.2477 | -3.6 | 3.7 |
| Dimethyl-lysine | STD | C8H18N2O2 | [M+H]+ | 175.1447 | 175.1437 | -5.3 | 22.2 |
| Dimethyl-arginine | STD | C8H18N4O2 | [M+H]+ | 203.1508 | 203.1499 | -4.3 | 22.7 |
| Dimethyl-guanosine | STD | C12H17N5O5 | [M+H]+ | 312.1308 | 312.1297 | -3.6 | 5.7 |
| Dimethyl-proline | STD | C7H13NO2 | [M+H]+ | 144.1025 | 144.1014 | -7.2 | 7.9 |
| Dimethyl-xanthine | STD | C7H8N4O2 | [M+H]+ | 181.0726 | 181.0716 | -5.5 | 4.2 |
| Diphosphoglycerate | STD | C3H8O10P2 | [M-H]- | 264.9520 | 264.9514 | -2.2 | 18.3 |
| Dodecanoyl-carnitine | STD | C19H37NO4 | [M+H]+ | 344.2801 | 344.2786 | -4.4 | 3.5 |
| Ergothioneine | STD | C9H15N3O2S | [M+H]+ | 230.0963 | 230.0954 | -4.0 | 13.6 |
| Fructose-1,6-diphosphate | STD | C6H14O12P2 | [M-H]- | 338.9888 | 338.9882 | -1.7 | 18.1 |
| Fructose-6-phosphate | STD | C6H13O9P | [M-H]- | 259.0224 | 259.0219 | -2.1 | 15.6 |
| GDP | STD | C10H15N5O11P2 | [M-H]- | 442.0171 | 442.0167 | -0.9 | 17.7 |
| GDP-glucose | STD | C16H25N5O16P2 | [M-H]- | 604.0699 | 604.0696 | -0.5 | 17.9 |
| Gluconate | STD | C6H12O7 | [M-H]- | 195.0510 | 195.0507 | -1.7 | 12.5 |
| Glucose-6-phosphate | STD | C6H13O9P | [M-H]- | 259.0224 | 259.0218 | -2.4 | 16.6 |
| Glutamate | STD | C5H9NO4 | [M-H]- | 146.0459 | 146.0457 | -1.1 | 13.6 |
| Glutamine | STD | C5H10N2O3 | [M+H]+ | 147.0770 | 147.0760 | -6.8 | 14.2 |
| Glutathione disulfide | STD | C20H32N6O12S2 | [M+H]+ | 613.1598 | 613.1581 | -2.8 | 17.3 |
| Glyceraldehyde-3-phosphate | STD | C3H7O6P | [M-H]- | 168.9907 | 168.9906 | -0.8 | 14.5 |
| Glycerate | STD | C3H6O4 | [M-H]- | 105.0193 | 105.0196 | 2.8 | 8.0 |
| Glycerol-phosphate | STD | C3H9O6P | [M-H]- | 171.0064 | 171.0062 | -1.4 | 14.2 |
| Glycerophosphocholine | STD | C8H20NO6P | [M+H]+ | 258.1106 | 258.1097 | -3.6 | 14.0 |
| Glycerophosphoethanolamine | MS/MS | C5H14NO6P | [M-H]- | 214.0486 | 214.0482 | -1.7 | 15.1 |
| Glycochenodeoxycholate | STD | C26H43NO5 | [M-H]- | 448.3068 | 448.3067 | -0.4 | 3.1 |
| GTP | STD | C10H16N5O14P3 | [M-H]- | 521.9834 | 521.9836 | 0.5 | 19.0 |
| Hexanoyl-carnitine | STD | C13H25NO4 | [M+H]+ | 260.1862 | 260.1853 | -3.6 | 4.6 |
| Hippurate | STD | C9H9NO3 | [M+H]+ | 180.0661 | 180.0651 | -5.2 | 3.6 |
| Histidine | STD | C6H9N3O2 | [M-H]- | 154.0622 | 154.0621 | -0.9 | 13.6 |
| Hypoxanthine | STD | C5H4N4O | [M+H]+ | 137.0463 | 137.0453 | -7.3 | 7.2 |
| Indoxyl-sulfate | STD | C8H7NO4S | [M-H]- | 212.0023 | 212.0019 | -1.8 | 4.3 |
| Isoleucine | STD | C6H13NO2 | [M-H]- | 130.0874 | 130.0874 | 0.3 | 9.0 |
| Keto(iso)leucine | STD | C6H10O3 | [M-H]- | 129.0557 | 129.0558 | 0.7 | 3.2 |
| Kynurenine | STD | C10H12N2O3 | [M+H]+ | 209.0926 | 209.0916 | -5.0 | 8.4 |
| Leucine | STD | C6H13NO2 | [M-H]- | 130.0874 | 130.0873 | -0.1 | 8.3 |
| Lysine | STD | C6H14N2O2 | [M+H]+ | 147.1134 | 147.1124 | -6.7 | 25.6 |
| Malate | STD | C4H6O5 | [M-H]- | 133.0142 | 133.0142 | -0.2 | 15.1 |

| | | | | | | | |
|---|---|---|---|---|---|---|---|
| Methionine | STD | C5H11NO2S | [M+H]+ | 150.0589 | 150.0579 | -6.7 | 9.3 |
| myo-Inositol | STD | C6H12O6 | [M-H]- | 179.0561 | 179.0558 | -1.6 | 16.3 |
| N-Acetyl-arginine | STD | C8H16O3N4 | [M+H]+ | 217.1301 | 217.1291 | -4.4 | 14.2 |
| N-Acetyl-aspartate | STD | C6H9NO5 | [M-H]- | 174.0408 | 174.0406 | -1.4 | 13.6 |
| N-Acetyl-glucosamine | STD | C8H15NO6 | [M+H]+ | 222.0978 | 222.0969 | -4.0 | 9.7 |
| N-Methyl-adenosine | STD | C11H15N5O4 | [M+H]+ | 282.1202 | 282.1179 | -8.3 | 12.2 |
| N1-Methyl-histidine | STD | C7H11N3O2 | [M+H]+ | 170.0930 | 170.0920 | -5.8 | 11.0 |
| N2-Acetyl-lysine | STD | C8H16O3N2 | [M+H]+ | 189.1239 | 189.1229 | -5.2 | 14.5 |
| N3-Methyl-histidine | STD | C7H11N3O2 | [M+H]+ | 170.0930 | 170.0919 | -5.9 | 12.2 |
| N6-Acetyl-lysine | STD | C8H16O3N2 | [M+H]+ | 189.1239 | 189.1229 | -5.4 | 11.9 |
| N6-Methyl-lysine | STD | C7H16N2O2 | [M+H]+ | 161.1290 | 161.1280 | -6.2 | 24.5 |
| NAD+ | STD | C21H27N7O14P2 | [M+H]+ | 664.1169 | 664.1147 | -3.3 | 13.7 |
| NADP+ | STD | C21H28N7O17P3 | [M+H]+ | 744.0833 | 744.0813 | -2.7 | 16.9 |
| Nicotinamide | STD | C6H6N2O | [M+H]+ | 123.0558 | 123.0548 | -8.3 | 4.7 |
| Octanoyl-carnitine | STD | C15H29NO4 | [M+H]+ | 288.2175 | 288.2165 | -3.6 | 3.8 |
| Ophthalmic acid | STD | C11H19N3O6 | [M+H]+ | 290.1352 | 290.1342 | -3.5 | 12.4 |
| Ornithine | STD | C5H12N2O2 | [M-H]- | 131.0826 | 131.0826 | 0.4 | 23.6 |
| Pantothenate | STD | C9H17NO5 | [M+H]+ | 220.1185 | 220.1176 | -4.1 | 4.8 |
| Pentose-phosphate | STD | C5H11O8P | [M-H]- | 229.0119 | 229.0115 | -1.5 | 15.3 |
| Phenylalanine | STD | C9H11NO2 | [M+H]+ | 166.0868 | 166.0858 | -6.2 | 7.2 |
| Phosphocholine | STD | C5H14NO4P | [M+H]+ | 184.0739 | 184.0729 | -5.1 | 15.0 |
| Phosphocreatine | STD | C4H10N3O5P | [M+H]+ | 212.0436 | 212.0428 | -4.0 | 14.7 |
| Phosphoenolpyruvate | STD | C3H5O6P | [M-H]- | 166.9751 | 166.9748 | -1.9 | 17.5 |
| Phosphoethanolamine | STD | C2H8NO4P | [M-H]- | 140.0118 | 140.0118 | -0.4 | 15.8 |
| Phosphoglycerate | STD | C3H7O7P | [M-H]- | 184.9857 | 184.9854 | -1.3 | 16.8 |
| Proline | STD | C5H9NO2 | [M+H]+ | 116.0712 | 116.0701 | -8.8 | 11.1 |
| Propionyl-carnitine | STD | C10H19NO4 | [M+H]+ | 218.1392 | 218.1383 | -4.3 | 7.0 |
| Pseudouridine | STD | C9H12N2O6 | [M-H]- | 243.0623 | 243.0617 | -2.4 | 9.7 |
| Quinolinic acid | STD | C7H5NO4 | [M-H]- | 166.0146 | 166.0143 | -1.7 | 14.0 |
| S-Adenosyl-homocysteine | STD | C14H20N6O5S | [M+H]+ | 385.1294 | 385.1283 | -2.9 | 13.0 |
| S-Adenosyl-methionine | STD | C15H22N6O5S | [M+H]+ | 399.1451 | 399.1437 | -3.3 | 17.0 |
| S-Methyl-ergothioneine | STD | C10H17N3O2S | [M+H]+ | 244.1120 | 244.1109 | -4.2 | 8.0 |
| Sedoheptulose-7-phosphate | STD | C7H15O10P | [M-H]- | 289.0330 | 289.0325 | -1.6 | 15.9 |
| Serine | STD | C3H7NO3 | [M+H]+ | 106.0504 | 106.0494 | -10.0 | 14.8 |
| Succinate | STD | C4H6O4 | [M-H]- | 117.0193 | 117.0195 | 1.7 | 14.1 |
| Taurine | STD | C2H7NO3S | [M+H]+ | 126.0225 | 126.0215 | -8.2 | 13.1 |
| Threonine | STD | C4H9NO3 | [M+H]+ | 120.0661 | 120.0650 | -8.6 | 13.1 |
| Trimethyl-histidine | STD | C9H15N3O2 | [M+H]+ | 198.1243 | 198.1233 | -4.9 | 10.7 |
| Trimethyl-lysine | STD | C9H20N2O2 | [M+H]+ | 189.1603 | 189.1594 | -4.9 | 23.2 |
| Trimethyl-phenylalanine | MS/MS | C12H17NO2 | [M+H]+ | 208.1338 | 208.1328 | -4.4 | 5.2 |
| Trimethyl-tryptophan | STD | C14H18N2O2 | [M+H]+ | 247.1447 | 247.1437 | -3.8 | 5.8 |
| Trimethyl-tyrosine | MS/MS | C12H17NO3 | [M+H]+ | 224.1287 | 224.1278 | -3.9 | 7.5 |
| Tryptophan | STD | C11H12N2O2 | [M+H]+ | 205.0977 | 205.0968 | -4.5 | 9.5 |
| Tyrosine | STD | C9H11NO3 | [M+H]+ | 182.0817 | 182.0808 | -5.3 | 11.8 |
| UDP | STD | C9H14N2O12P2 | [M-H]- | 402.9949 | 402.9945 | -1.1 | 16.3 |
| UDP-glucose | STD | C15H24N2O17P2 | [M-H]- | 565.0477 | 565.0473 | -0.7 | 16.0 |
| UDP-glucuronate | STD | C15H22N2O18P2 | [M-H]- | 579.0270 | 579.0266 | -0.7 | 18.6 |
| UDP-N-acetyl-glucosamine | STD | C17H27N3O17P2 | [M-H]- | 606.0743 | 606.0738 | -0.8 | 14.7 |
| UMP | STD | C9H13N2O9P | [M-H]- | 323.0286 | 323.0281 | -1.5 | 14.4 |
| Urate | STD | C5H4N4O3 | [M+H]+ | 169.0362 | 169.0351 | -6.2 | 12.0 |
| Uridine | STD | C9H12N2O6 | [M-H]- | 243.0623 | 243.0612 | -4.5 | 6.9 |
| UTP | STD | C9H15N2O15P3 | [M-H]- | 482.9613 | 482.9610 | -0.5 | 17.6 |
| (iso)Valeryl-carnitine | STD | C12H23NO4 | [M+H]+ | 246.1705 | 246.1696 | -3.6 | 5.2 |
| Valine | STD | C5H11NO2 | [M-H]- | 116.0717 | 116.0719 | 1.7 | 10.7 |
| Xanthine | STD | C5H4N4O2 | [M-H]- | 151.0261 | 151.0260 | -0.8 | 8.0 |
